# *Ab initio* modelling of an essential mammalian protein: Transcription Termination Factor 1 (TTF1)

**DOI:** 10.1101/2021.11.11.468186

**Authors:** Kumud Tiwari, Aditi Gangopadhyay, Gajender Singh, Samarendra Kumar Singh

## Abstract

Transcription Termination Factor 1 (TTF1) is an essential mammalian protein that regulates cellular transcription, replication fork arrest, DNA damage repair, chromatin remodelling etc. TTF1 interacts with numerous cellular proteins to regulate various cellular phenomena, and plays a crucial role in maintaining normal cellular physiology, dysregulation of which has been reported towards cancerous transformation of the cells. However, despite its key role in cellular physiology, the complete structure of human TTF1 has not been elucidated to date, either experimentally or computationally. Hence, understanding the structure of human TTF1 becomes highly important for studying its functions and interactions with other cellular factors. Therefore, the aim of this study was to construct the complete structure of human TTF1 protein, using molecular modelling approaches. Owing to the lack of suitable homologues in the PDB, the complete structure of human TTF1 was constructed using *ab initio* modelling. The structural stability was determined using molecular dynamics (MD) simulations in explicit solvent, and trajectory analyses. The representative structure of human TTF1 was obtained by trajectory clustering, and the central residues were determined by centrality analyses of the residue interaction network of TTF1. Two residue clusters, in the oligomerisation domain and C-terminal domain, were determined to be central to the structural stability of human TTF1. To the best of our knowledge, this study is the first to report the complete structure of human TTF1, and the results obtained herein will provide structural insights for future research in cancer biology and related studies.

**Author Summary:** The transcription termination factor 1 (TTF1) is an essential multifunctional mammalian protein which plays important role in regulating important cellular process like transcription, replication, DNA damage repair, chromatin remodelling etc. and its dysregulation leads to various cancers. Despite its being such an important factor, the complete structure of human TTF1 has not been determined to date, either using experimental techniques or computationally. Therefore, the aim of this study was to construct the complete structure of human TTF1 using computational modelling. In this study the complete structure of human TTF1 was constructed by *ab initio* modelling using iTasser. The stability of this model was determined by 200 ns molecular dynamics (MD) simulations. The representative conformation of human TTF1 was further determined by clustering the simulation trajectory and the residues that are central to the stability of this structure were identified. The results demonstrate the presence of two residue clusters in human TTF1, one in the oligomerisation domain and other in the C-terminal domain, which were found to be crucial for the structural stability of this protein. Hence, the results of this study will aid future studies in this field towards engineering this important protein for further biochemistry and cell biology research.

## 1. Introduction

Ribosomes are essential cellular organelles that partake in protein synthesis in both prokaryotes and eukaryotes. Ribosomes are comprised of ribosomal proteins and ribosomal RNA (rRNA), which is encoded by ribosomal DNA (rDNA), and serves as the catalytic subunit of the protein translation machinery. Eukaryotic rDNA is distributed in clusters of ∼300-400 copies at both ends of the respective chromosomes (acrocentric chromosomes: 13, 14, 15, 21, and 22). These tandem repeats of rDNA copies create dense chromosomal regions called **N**ucleolar **O**rganizer **R**egions (**NOR**s) which consists of a non-transcribed spacer region flanked by pre-RNA coding regions. Of the total RNA that is transcribed, 80% consists of rRNAs [1,2]. Both the initiation and termination of rDNA transcription is mediated by a transcriptional regulator called Transcription Termination Factor 1 (TTF1), which is an essential protein in mammalian cells. The gene encoding TTF1 is located on **9q34**.**13**, in the long arm of chromosome 9. Transcription Termination Factor 1 protein (TTF1p) binds to DNA elements known as Sal box, located upstream and downstream of the rDNA gene repeats. In mammalian cells, the Sal box element consists of a *Sal*I restriction site within the 11 bp sequence, G***GGTCGACC***AG [3]. Following its discovery as a transcription regulator, subsequent studies demonstrated that TTF1 is involved in polar replication fork arrest and also acts as a chromatin remodelling factor [4,5]. Current findings demonstrate that TTF1p interacts with various DNA damage sensing proteins, including Cockayne Syndrome B (CSB) [6], Mouse Double Minute 2 (MDM2) [7] and tumor suppressor Alternative Reading Frame (ARF) [8] protein, but the mechanism and exact roles of TTF1p remains to be identified to date. The overexpression of TTF1 has been corelated in various tumours, which indicates that owing to tumor hyperproliferation, TTF1 is required in higher quantities to meet the higher rate of ribosome biogenesis in tumor cells [9–11]. The TTF1 protein has several other unidentified roles, as it appears to interact with various other factors necessary for regulating a wide variety of physiological phenomena in cells. TTF1 is truly a multifunctional protein, and hence, it becomes important to characterise the numerous unidentified roles of this protein in cellular physiology both in healthy and cancerous cells.

TTF1p has distinct functional domains, including an N-terminal regulatory domain (NRD), which also is responsible for the oligomerisation of TTF1 [12]. It has been shown that due to its oligomerisation property, TTF1p can loop the ends of rDNA together, thereby placing the promoter and terminator regions in proximity to efficiently recycle the transcription machinery, and this model is known as the “ribomotor” model [13]. Furthermore, TTF1 has a functional central domain and a C-terminal domain, which is essential for the activation and termination of Pol I-mediated transcription on a nucleosomal rDNA template [14]. The central domain has the highly conserved DNA binding myb/SANT-like domain which has strong homology with the DNA binding domain of Reb1 protein of *Schizosaccharomyces pombe*, and proto-oncoprotein c-Myb [15,16].

The only crystal structure of its yeast homolog protein, RNA Polymerase I enhancer binding protein (Reb1p) [17], bound to DNA, was solved to atomic resolution by our group [15]. The structure clearly shows an N-terminal regulatory domain which is also known as the dimerization domain, a central DNA-binding domain, and the C-terminal transcriptional terminator domain. Using various mutants, it was demonstrated that the mere binding of DNA to Reb1p is not sufficient for terminating transcription. Further it was shown that the interaction of Reb1p with Replication Protein A (RPA), via the C-terminal domain of Reb1p, is an essential requirement for effective transcriptional termination. The interaction with RPA induces an allosteric change which is necessary for stopping the movement of RNA polymerase I. Also, the domain of Reb1p which binds to DNA was identified to atomic resolution and the residues involved in protein-DNA contacts were identified. This region consists of two myb-associated domains (mybAD1 and mybAD2) and two Myb repeats (mybR1 and mybR2). The helices involved in this region make contact with DNA at various residues [17].

TTF1 is an essential cellular protein owing to its numerous roles in several vital cellular functions, which are necessary for maintaining healthy cellular physiology. Understanding the structure of TTF1 would provide insights into the mechanistic aspect of its function. To date, there are no experimentally-determined structures or *in silico* models of TTF1. Our lab is involved in purifying and physically solving the structure of this protein. So far, crystallization trials have proved to be unsuccessful, and we are therefore attempting cryo-EM studies as well. Alternatively, computational modelling studies on this essential protein will provide a better understanding so that we can engineer the protein for future studies.

In the absence of experimentally-derived structures, homology modelling serves as a reliable method for the construction of protein structures. However, the reliability of the protein model depends on various factors, including the sequence identity between the template and target proteins. When the template-target identity falls below 30%, known as the twilight zone, the protein structure needs to be constructed by threading or *ab initio* methods [18]. This is due to the fact that below the twilight zone, the evolutionary relatedness between the template and target is doubtful, and the confidence of the prediction is low [19]. The worldwide experiment for protein structure prediction, Critical Assessment of protein Structure Prediction (CASP), ranked the iTasser (iterative threading assembly refinement) server as the best tool for *ab initio* protein modelling. In the latest CASP14 experiment conducted in 2020, the iTasser server (Zhang server) ranked the best among 47 groups [20,21]. The iTasser server also ranked best in the previous CASP7, CASP8, CASP9, CASP10, CASP11, CASP12, and CASP13 experiments [22]. In the CASP9 experiments in 2010, the iTasser server was predicted to the best tool for protein function prediction [21]. In this study, the structure of the TTF1 protein was constructed by molecular modelling, using the iTasser server. The predicted models were validated and the structure was subjected to molecular dynamics (MD) simulations for 200 ns for studying the structural stability of TTF1, and determining the most stable conformation of the protein. Our study aimed to predict the structure of TTF1, which is an essential protein, using computational modelling. The results of our study will prove to be important for understanding the structural, functional, and therapeutic role of this essential protein.

## 2. Results

### 2.1 Sequence-based analyses

The results of sequence-based analysis with ProtParam showed that TTF1 is an unstable hydrophilic protein, as revealed by an instability index of 51.13 and grand average of hydrophobicity (GRAVY) of -0.939. This was corroborated by the results of disorder prediction, which showed that more than 50% of the residues of TTF1 are disordered (Fig 1). The results of disorder prediction further revealed that residues 1-3, 689-696, 700-701, 709, and 903-905 were disordered and had protein binding properties (S1 Fig). The physicochemical properties predicted by ProtParam and anticipation of disulphide bond (S-S) pattern by CYS REC tool are enlisted in Table 1.

**Table 1:** Physicochemical properties of TTF1, as determined with ProtParam.

| Physicochemical Properties | Values |
| --- | --- |
| Number of residues | 905 |
| Molecular formula | C<sub>4512</sub>H<sub>7282</sub>N<sub>1302</sub>O<sub>1399</sub>S<sub>28</sub> |
| Molecular weight | 103 kDa |
| Theoretical pI | 9.41 |
| Instability index | 51.13 |
| Aliphatic index | 65.09 |
| Total number of negatively charged residues (D+E) | 135 |
| Total number of positively charged residues (R+K) | 171 |
| Extinction coefficient | 96760 |
| Grand average of hydrophobicity (GRAVY) | -0.939 |
| Estimated half-life (mammalian reticulocyte, *in vitro*) | 30 hours |
| S-S Bond (predicted) | 22-737, 73-887, 445-892 |

**Fig 1:**
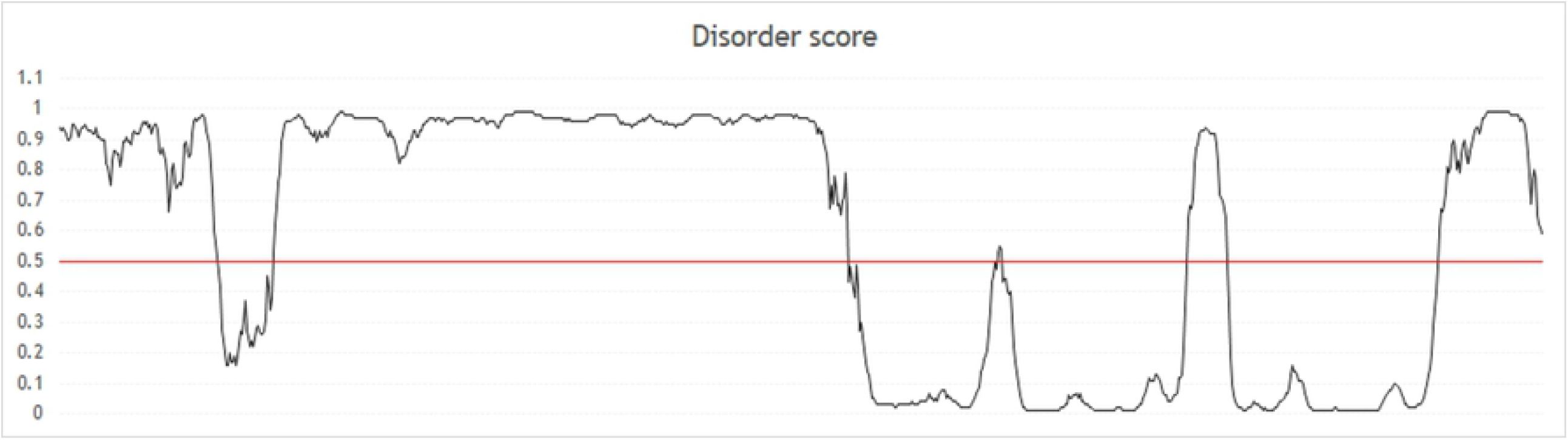
Graphical representation of the disordered regions of the human TTF1 protein. Residues with disorder score ≥ 0.5 (represented by the horizontal red line) were considered to be disordered.

### 2.2 Ab inito modelling and structural validation of TTF1

The results of template search using BLASTp against the PDB revealed that the highest target-template coverage was 4%, which was well below the twilight zone for homology modelling [23]. Therefore, the structure of human TTF1 could not be modelled using the template-based methods in comparative modelling. The complete structure of human TTF1p was therefore modelled using *ab initio* methods, using the iTasser server. The confidence of the models predicted by iTasser are indicated by the C-score, which is a confidence score that provides a measure of the quality of the models generated by iTasser. The C-scores range between -5 and 2, with higher values indicating predictions of higher confidence, while lower values of C-score indicate predictions of lower confidence [20]. In this study, the model with the highest C-score of -0.60 was selected for subsequent analyses. This model was further minimised using Yasara, and the energy minimised structure was validated using ProSA [24,25]. The results of ProSA validation revealed that the structure of TTF1 was comparable to structures of similar size in the PDB, which had been determined using X-ray crystallography (Fig 2A). Analysis of the Ramachandran plot with Procheck revealed that only 1.0% of the residues were in the disallowed regions of the plot, while 82.9% and 14.1% of the residues were in the most favoured and additional allowed regions, respectively (Fig 2B).

**Fig 2:**
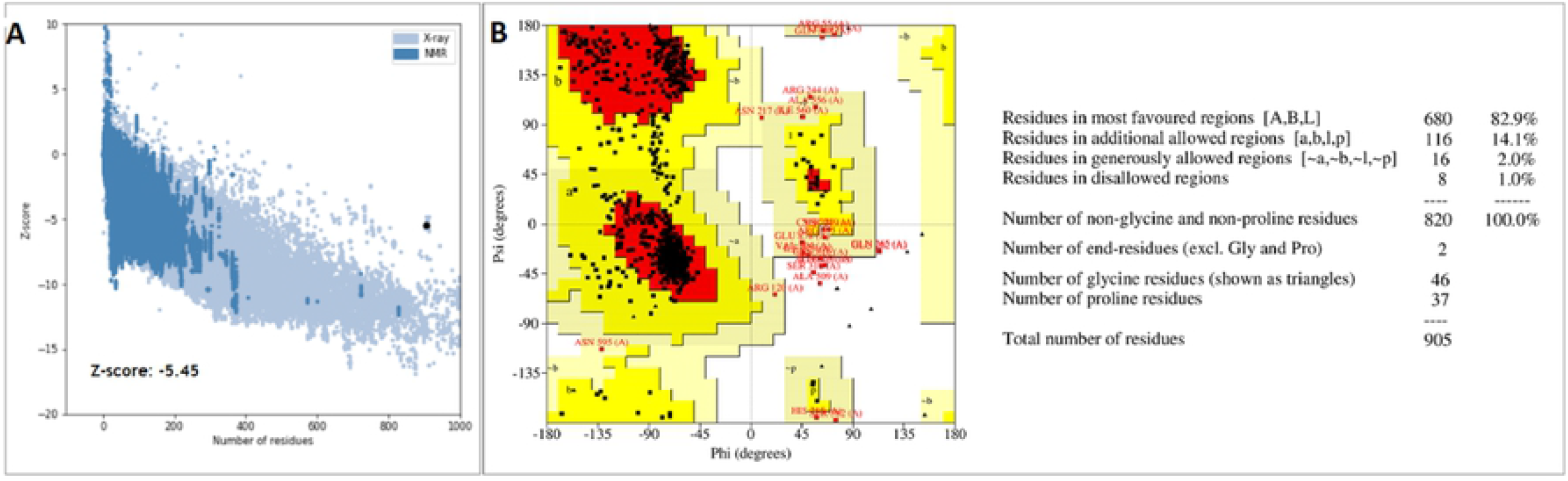
Structural validation of the energy minimised model of TTF1 using **A)** ProSA and **B)** Ramachandran plot analysis with Procheck.

### 2.3. Functional validation of TTF1

The results of analysis with TM-align revealed that the model of TTF1 generated by iTasser (Fig 3) was structurally most similar to cas13b (PDB ID: 6AAY), which is an RNA-binding protein from *Bergeyella zoohelcum* with RNase activity [26]. The human TTF1 protein is a DNA-binding protein that plays an important role in transcriptional termination. The TM-score of the alignment was 0.960, indicating correct topology, and the RMSD between the generated model of TTF1 and cas13b was 2.29 Å, indicating high structural similarity between the two proteins. The structural similarity between TTF1 and cas13b indicated that the model of TTF1 obtained herein, possesses potential nucleic acid binding properties, similar to cas13b.

**Fig 3:**
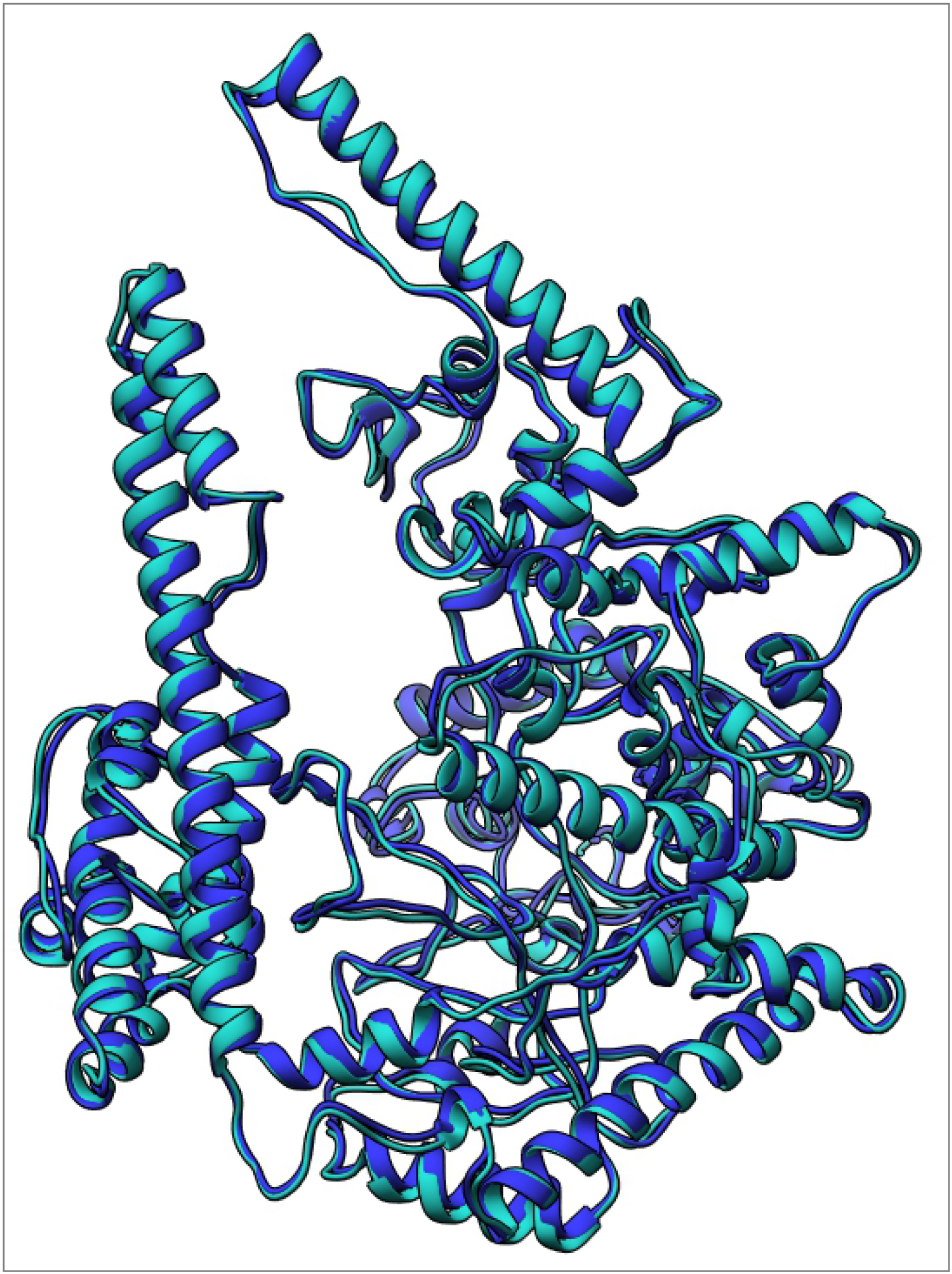
The structure of TTF1 constructed by *ab initio* modelling is depicted in light blue ribbon representation, and the structure obtained after minimisation with Yasara is depicted in dark blue ribbon representation.

The results of ligand binding analyses with COFACTOR and COACH revealed that residues 620-626 of the TTF1 model have potential binding property to the ligand phosphoaminophosphonic acid-adenylate ester (ANP). ANP is a non-hydrolysable analogue of ATP, and comprises triphosphate, adenine, and ribose sugar moieties, similar to the composition of DNA. This indicated that the model of TTF1 predicted using iTasser has potential nucleic acid binding properties, and logically relates to the DNA-binding properties of TTF1 reported in literature and also been validated in our lab using the purified TTF1 protein [3]. The ligand binding properties of the model of TTF1 were predicted to be most similar to those of the recombinase A protein of *Escherichia coli* (PDB ID: 3CMV), which possesses single-stranded DNA binding properties. These results indicated that the model of TTF1 possesses potential DNA binding properties, in agreement with the reports in existing literature and our experimental data (not shown here). The results of CD analyses revealed that residues 621-677 of TTF1 comprises the Myb-like DNA binding domain of TTF1 (pfam accession number: 13921) (S 2 Fig). The results of sequence-based CD analysis of TTF1 corroborated with the results of structure-based ligand binding site prediction by COFACTOR and COACH, further validating the DNA-binding potential, and thus the functional potential of the model of TTF1 constructed using iTasser. Furthermore, the residues with ANP-binding properties mapped to the DNA-binding domain of TTF1, implying the potential nucleotide binding properties of the structure of TTF1 generated by iTasser.

The results of consensus-based GO prediction revealed that the molecular function of the TTF1 protein model was associated with GO terms GO:0035639 (purine ribonucleoside triphosphate binding), GO:0032559 (adenyl ribonucleotide binding), and GO:0043167 (ion binding), with GO scores of 0.40, 0.40, and 0.39, respectively. These results further confirmed the nucleotide binding properties of the structure of TTF1 obtained with iTasser. The results of functional validation thus implied that the TTF1 model obtained using *ab initio* modelling has potential nucleic acid-binding properties, and agrees with the data reported in literature and observed in our lab.

### 2.4. Trajectory analyses

The structural model of TTF1 thus obtained by *ab initio* modelling was subjected to 200 ns MD simulations for investigating the structural stability and determining any possible conformational changes in TTF1. Trajectory visualisation revealed that the protein stabilised after 20 ns and remained stable thereafter. This was further observed in the values of root mean square deviation RMSD, which became steady after 20 ns (Fig 4). As depicted in the Fig 4A, the values of RMSD became increasingly steady after 100 ns, and remained steady thereafter, with fluctuations in the RMSD values being in the range of 1-1.5 Å. This indicated that the system had reached equilibrium after 100 ns and remained stable thereafter. This was further corroborated by the values of RoG (Fig 4B), which remained steady after 100 ns. The RoG is an indicator of structural compactness, and fluctuations in the values of RoG indicate protein unfolding. The fact that the values of RoG became steady after 100 ns indicated that the structure of TTF1 was stable and compact during the production run. Analysis of the values of RMSF revealed that some residues had higher flexibility, as indicated by the RMSF values, which were higher than 1.5 Å. The higher flexibility of these residues could be attributed to the fact that these residues mapped to the disordered regions predicted using DisoPred (Fig 4C).

**Fig 4:**
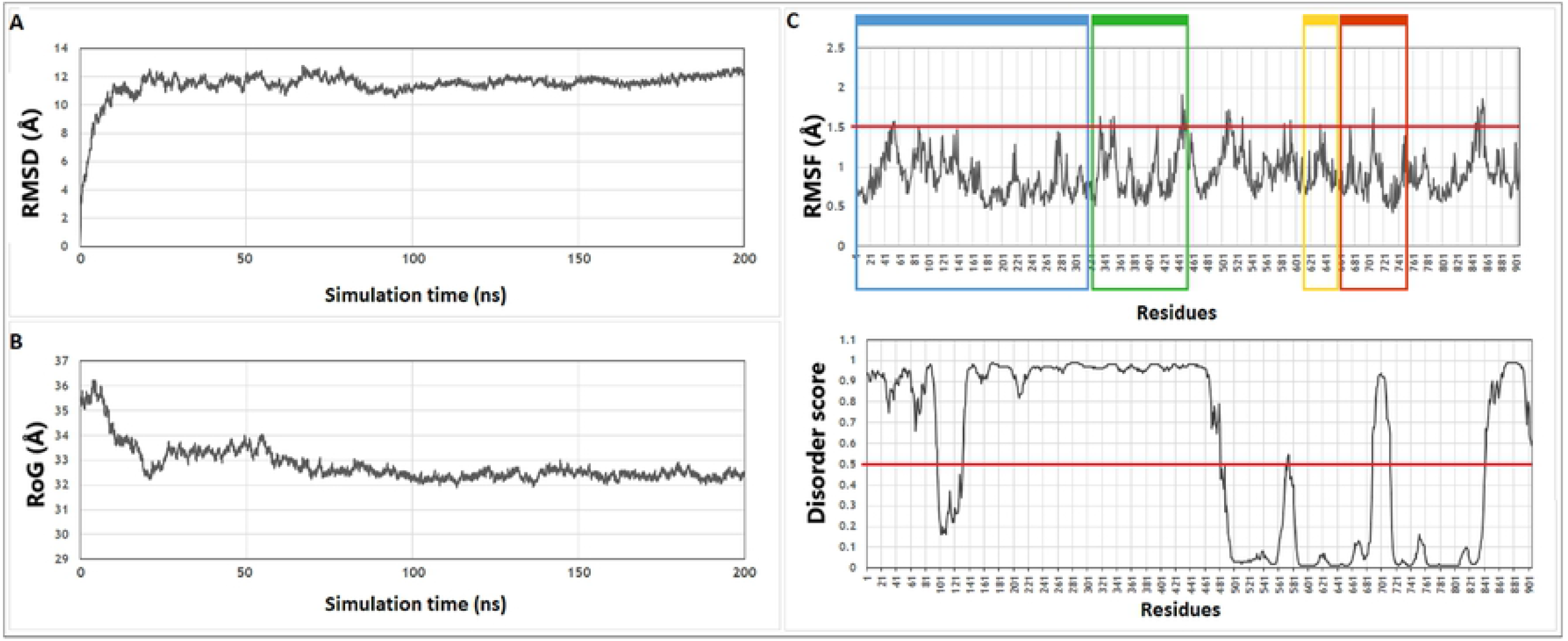
Graphical representation of the values of **A)** RMSD and **B)** RoG of the protein backbone throughout the trajectory. **C)** Comparison of the average RMSF values and disorder scores of TTF1. The oligomerisation domain (residues 1-320), Myb domain 1 (residues 612-660), Myb domain 2 (residues 661-745), and chromatin remodelling region (residues 323-445) are indicated by blue, yellow, red, and green rectangles, respectively.

### 2.5. Representative structure of TTF1

The trajectory was clustered using Chimera v1.14, and the representative frame of the most populated cluster was selected as the representative conformation of TTF1 (Fig 5A, refer to the supplementary for coordinate file). The oligomerisation domain (residues 1-320), Myb domain 1 (residues 612-660), Myb domain 2 (residues 661-745), and chromatin remodelling region (residues 323-445,) [5,12,16] were mapped to the complete structure of TTF1 obtained herein (Fig 5B).

**Fig 5:**
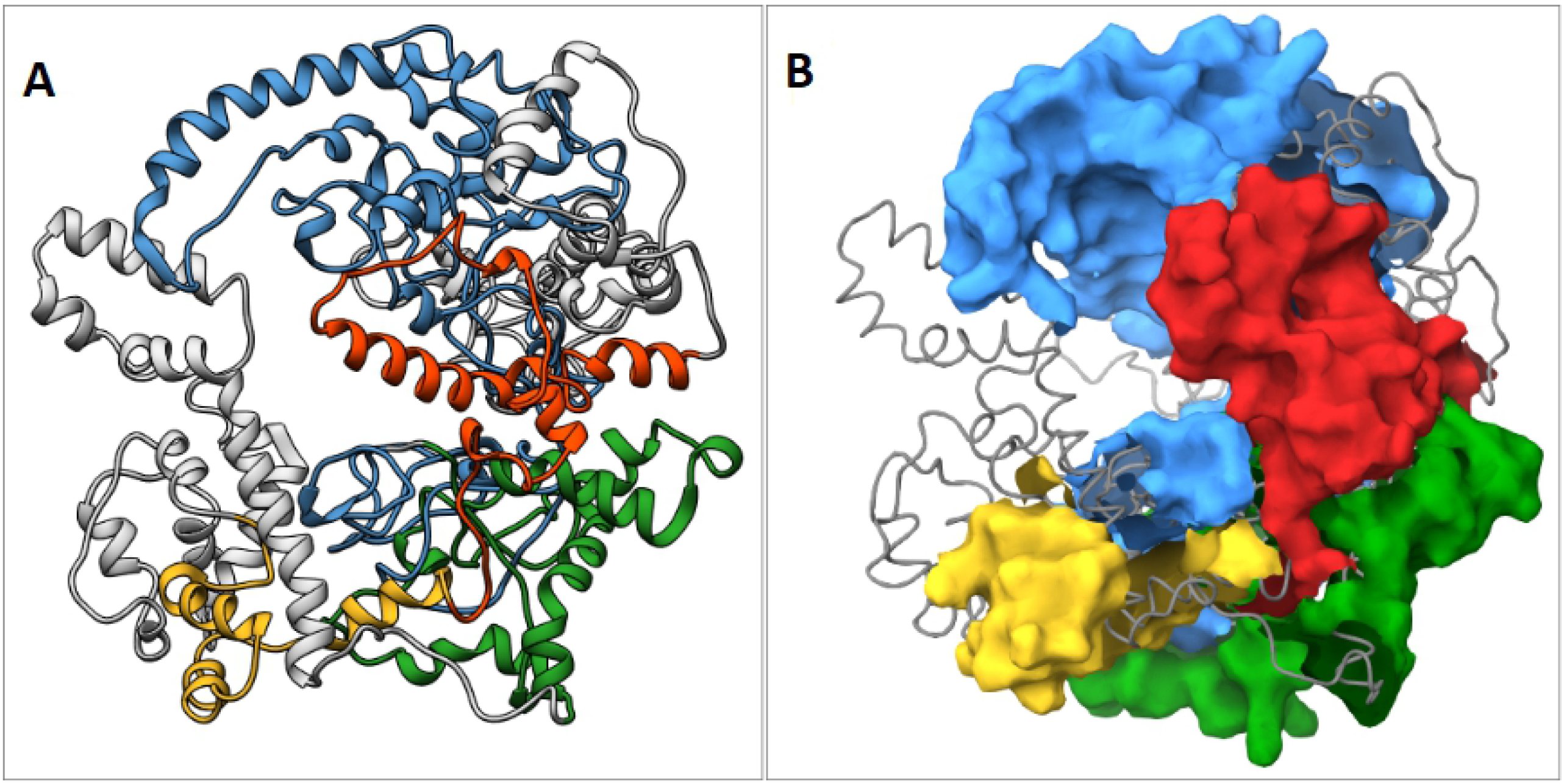
**A)** Ribbon and **B)** Surface representation of the representative structure of TTF1 obtained by trajectory clustering. The oligomerisation domain, Myb domain 1, Myb domain 2, and chromatin remodelling region are represented in blue, yellow, red, and green, respectively.

### 2.6. Centrality analyses

The RINs of the representative structure of TTF1 was determined using Cytoscape v3.8.2 (Fig 6), and the central residues were identified using the RINspector plugin, based on the RCA Z-scores. In the RIN, the nodes indicate the residues, while the edges represent the intra-residue interactions. Residues with RCA Z-scores ≥ 2 were considered to be central to the structural stability of the protein. As depicted in Fig 6, the residues with Z-scores ≥ 2 are coloured in yellow, and those with Z-scores ≥ 2 are represented in red. The bigger nodes indicate residues with higher values of Z-scores. The RIN revealed two interaction clusters, with one cluster being located in the oligomerisation domain of TTF1, and the other being located towards the C-terminal region of the protein (Fig 6A and 6B). The Z-scores of the residues in the interaction cluster in the oligomerisation domain were higher than those of the residues in the C-terminal domain, indicating that the interaction cluster in the oligomerisation domain plays a more crucial role in the stability of the human TTF1 protein than that of the interaction cluster in the C-terminal domain. The Z-scores of the central residues determined by centrality analysis are enlisted in Table 2.

**Table 2:** Central residues of TTF1, determined by centrality analysis.

| Central residue | Corresponding domain | RCA Z-score | Secondary structure |
| --- | --- | --- | --- |
| K68 | Oligomerisation | 7.246 | Loop |
| E196 | Oligomerisation | 7.2 | Helix |
| R71 | Oligomerisation | 7.161 | Loop |
| E195 | Oligomerisation | 7.111 | Loop |
| Q30 | Oligomerisation | 7.097 | Helix |
| HIE34 | Oligomerisation | 6.991 | Helix |
| W198 | Oligomerisation | 6.813 | Helix |
| R38 | Oligomerisation | 6.682 | Helix |
| G202 | Oligomerisation | 6.555 | Loop |
| E35 | Oligomerisation | 6.453 | Helix |
| K17 | Oligomerisation | 6.366 | Helix |
| E206 | Oligomerisation | 5.788 | Helix |
| K31 | Oligomerisation | 4.573 | Helix |
| S226 | Oligomerisation | 4.417 | Loop |
| E27 | Oligomerisation | 4.309 | Helix |
| T186 | Oligomerisation | 4.204 | Loop |
| N228 | Oligomerisation | 3.947 | Loop |
| R229 | Oligomerisation | 3.614 | Loop |
| S23 | Oligomerisation | 3.159 | Helix |
| I660 | myb/SANT-like 1 | 2.83 | Helix |
| R664 | myb/SANT-like 2 | 2.825 | Loop |
| K434 | Chromatin remodelling | 2.782 | Helix |
| F657 | myb/SANT-like 1 | 2.778 | Helix |
| R164 | Oligomerisation | 2.769 | Loop |
| E165 | Oligomerisation | 2.693 | Loop |
| S895 | - | 2.693 | Helix |
| N893 | - | 2.661 | Loop |
| E882 | - | 2.608 | Loop |
| T896 | - | 2.596 | Helix |
| S161 | Oligomerisation | 2.594 | Loop |
| Q172 | Oligomerisation | 2.548 | Helix |
| E191 | Oligomerisation | 2.535 | Loop |
| E431 | Chromatin remodelling | 2.527 | Helix |
| G899 | - | 2.499 | Loop |
| A176 | Oligomerisation | 2.469 | Helix |
| S435 | Chromatin remodelling | 2.388 | Helix |
| R902 | - | 2.326 | Loop |
| R182 | Oligomerisation | 2.302 | Helix |
| D471 | - | 2.069 | Helix |

**Fig 6:**
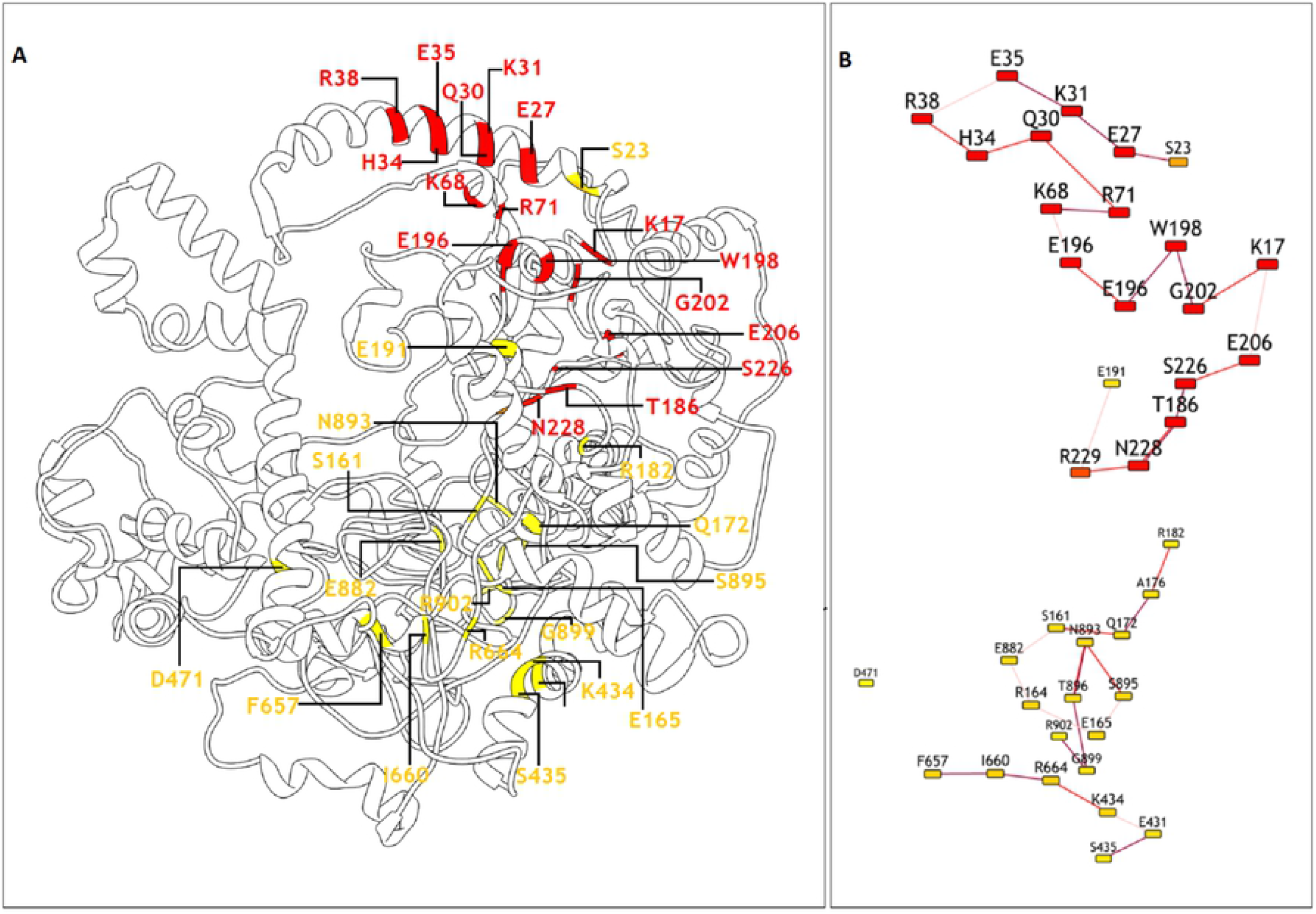
The central residues of TTF1 identified by RIN and centrality analyses in the **A)** 3-dimensional structure of TTF1. **B)** The central residues in the RIN of TTF1 in 2D representation. The nodes and edges represent the residues and inter-residue interactions, respectively. The size of the nodes corresponds to the value of the RCA Z-score, with bigger nodes corresponding to residues with higher values of Z-scores. The residues with RCA Z-score ≥ 2 and ≥ 4 are indicated in yellow and red, respectively.

### 2.7. Intra-residue hydrogen bonds

Hydrogen bonds with occupancy ≥ 75% and ≥ 85% throughout the 200 ns trajectory and in the last 50 ns, respectively, were considered to be important for the structural stability of the protein. The frequency of the hydrogen bonds throughout the trajectory and in the last 50 ns was determined using VMD. The occupancy of the intra-residue hydrogen bonds formed by the central residues is provided in S 1 Table, and the occupancy of all the intra-residue hydrogen bonds with occupancy ≥ 75% and ≥ 85% throughout the 200 ns trajectory and in the last 50 ns, respectively, are provided in the S 1 Table. The results of interaction analyses revealed that residues K17, E27, Q30, E35, R164, W198, and N228 of the oligomerisation domain, K434 of the chromatin remodelling region, and F657 of the myb/SANT-like-1 domain were most crucial to the structural stability of the protein, as indicated by the number of intra-residue hydrogen bonds and the occupancy of the hydrogen bonds throughout the trajectory.

## 3. Discussion

TTF1 is a crucial multifunctional nucleolar protein that regulates both transcription initiation as well as transcriptional termination of ribosomal genes by binding to specific motif sequence and also arrests of the replication fork in polar fashion [2]. In addition, TTF1 regulates the transcription of genes transcribed by RNA polymerase I. Using truncated human and murine TTF1 proteins, Evers and Grummt first reported species-specific sequence differences in the DNA-binding region of mammalian TTF1 [3]. Despite its major regulatory role in mammalian transcription, replication and chromatin remodelling, the complete structure of human TTF1 remains to be elucidated to date. A partial structure of human TTF1 has been predicted by AlphaFold v2.0, which uses artificial intelligence for predicting the 3-dimensional structure of proteins. However, the structure predicted by AlphaFold is partial (residues 491-866), and the remaining residues are largely unfolded, and the confidence of prediction of these unfolded regions is very low [27]. As all the residues of a protein are important for its complete regulation and function, it is necessary to consider that protein in its entirety in structural analyses. In this study, we therefore attempted to construct the complete structure of the human TTF1 protein using *ab initio* modelling and MD simulations, and also identified the residues that are central to the structural stability of human TTF1 by network analyses. To the best of our knowledge, this study is the first to report the complete structure of the human TTF1 protein (refer supplementary for coordinate file).

Owing to the lack of suitable structural homologues in the PDB with sequence coverage above the twilight zone, the structure of TTF1 was modelled using *ab initio* methods. The model of TTF1 thus obtained was subjected to functional validation and GO analysis for establishing the functional relevance. MD simulations are frequently used for obtaining atom-level insights into the structural dynamics and behaviour of biomolecular system. The stability of the model was subsequently evaluated by MD simulation for 200 ns, using an explicit TIP4P solvent, and the trajectory was analysed for investigating structural stability and hydrogen bond frequency. The representative conformation of the human TTF1 protein was obtained by trajectory clustering, and the residues that play a central role in the structural stability of TTF1 were identified by network analysis and determination of residue centrality. The results of RIN analysis and computation of centrality measures revealed two interaction clusters in the structure of human TTF1, with one in the oligomerisation domain of TTF1 and the other in the C-terminal domain. The data further indicated that the residue cluster in the oligomerisation domain plays a more significant role in the stability of TTF1, compared to that in the C-terminal domain. The N-terminal oligomerization domain has been shown to play important regulatory function [2] while the C-terminal domain is involved in transcription termination [5]. In the absence of experimentally-derived structural data pertaining to the human TTF1 protein, we believe that the results of our study provide valuable structural information, including domain architecture, and their characteristics, among others. Hence, our study could facilitate future studies aimed towards understanding the mechanism underlying the function of the human TTF1, including its interaction with other protein, and for engineering this protein with the purpose of solving its physical structure, drug design and therapeutic applications etc.

## 4. Conclusion

Conclusively, this is very first study to report complete structure of the essential human TTF1 protein, using computational modelling, and identify the residues and its characteristics that are central to the structural stability of the protein.

## 5. Materials and Methods

### 5.1 Sequence retrieval and sequence-based analyses

The sequence of TTF1 was retrieved from UniProtKB (UniProtKB accession number: Q15361). The physicochemical properties of TTF1 were analyzed using ProtParam [28], and the disorder profile was analyzed using DisoPred version 3.1 [29,30].

### 5.2 Ab initio modelling of TTF1

The structural homologues of human TTF1 in the PDB was searched using BLASTp and threading-based approaches, for identifying suitable templates for homology modelling. Owing to the lack of suitable structural templates, the structure of human TTF1 was modelled using *ab initio* modelling, using the iTasser server [20]. In the iTasser algorithm, the final models are selected using the SPICKER program for clustering the generated structures. The structure of TTF1 generated by iTasser was initially minimised using the Yasara energy minimization server, with the Yasara force field [24]. The energy minimised structure was then validated using Ramachandran plot analysis and ProSA [31,32].

### 5.3 Functional validation of TTF1 constructed by *ab initio* modelling

The models generated by iTasser were functionally validated using the TM-align program for determining the structures in the PDB that are structurally, and thus functionally, similar to the models of TTF1p constructed by *ab initio* modelling. The TM-align program was used to identify structures in the PDB that are structurally similar to the model generated by iTasser. This program determines the similarity between proteins on the basis of the TM-score, a scoring function that provides a quantitative measure of topological similarity between proteins [33]. It provides a measure of structural similarity, with values > 0.5 indicating models of correct topology [34]. The models were further validated using the COACH and COFACTOR programs for predicting the ligand binding sites, based on the similarity of the protein folds with functional templates [35,36]. The result of ligand binding site prediction was mapped to the results of sequence-based conserved domain (CD) analyses using the CD search tool of NCBI [37]. The molecular function of the modelled protein was further validated by consensus-based gene ontology (GO) search.

### 5.4 MD simulations

The model of TTF1 obtained by *ab initio* modelling was subjected to MD simulations for 200 ns using Flare v4, which is based on the OpenMM Toolkit, for studying the structural stability and determining any possible conformational changes of TTF1p. The protein was then prepared in Flare v4 at pH 7.4, and solvated in TIP4P solvent using a buffer of 10 Å thickness. The system was subsequently neutralised by the addition of 28 Cl^-^ ions. The system was then minimized until the energy tolerance reached 0.25 Kcal/mol, and subsequently equilibrated for 200 ps. It was then finally subjected to 200 ns MD simulations at a temperature of 298 K and a pressure of 1 bar, using the XED force field and the NPT ensemble. The timestep was set to 2 fs.

The values of root mean square deviation (RMSD), root mean square fluctuations (RMSF), and radius of gyration (RoG) of the protein backbone throughout the trajectory was analyzed using the vmdICE plugin in VMD v1.9.3 [38,39]. The occupancy of the inter-residue hydrogen bonds throughout the 200 ns trajectory and in the last 50 ns was determined using VMD v1.9.3. The portion of the trajectory following equilibration was clustered using Chimera v1.14, and the representative frame of the most populated cluster was selected as the representative conformation of TTF1p [40].

### 5.6 Determination of residue interaction networks (RINs) and centrality analysis

The RINs of TTF1p were determined using the RINalyzer plugin in Cytoscape v3.8.2 [41,42]. The network centrality measures were computed using the RINspector plugin in Cytoscape v3.8.2, based on the residue centrality analysis (RCA) Z-score.

## Acknowledgement

The authors are thankful to the Director Prof. A.K. Tripathi and Coordinator Prof. S.M. Singh, School of Biotechnology, Institute of Science, Banaras Hindu University for providing space and facilities to conduct the research. We thank Dr. V.K. Singh for providing valuable suggestions during the study. We are thankful to Department of Biotechnology, Govt. of India for funding Samarendra K Singh (SKS) and Kumud Tiwari (KT) with grant and fellowship respectively. Author Aditi Gangopadhyay (AG) acknowledges the Council of Scientific and Industrial Research (CSIR), New Delhi, for providing financial assistance. We also thank CSIR for funding the fellowships of Gajender Singh (GS).

## Author contribution

SKS was involved in the Conceptualization and designing; Supervision; Writing - critical review & editing of the manuscript. KT and AG involved in Data curation; Formal analyses; Investigation; Methodology; Project administration; Resources; Software; Validation; Visualization; Writing - original draft; review & editing of the manuscript. GS contributed analyses tools and data; Visualization; Writing – original draft.

## Supporting information

**S 1 Fig.**
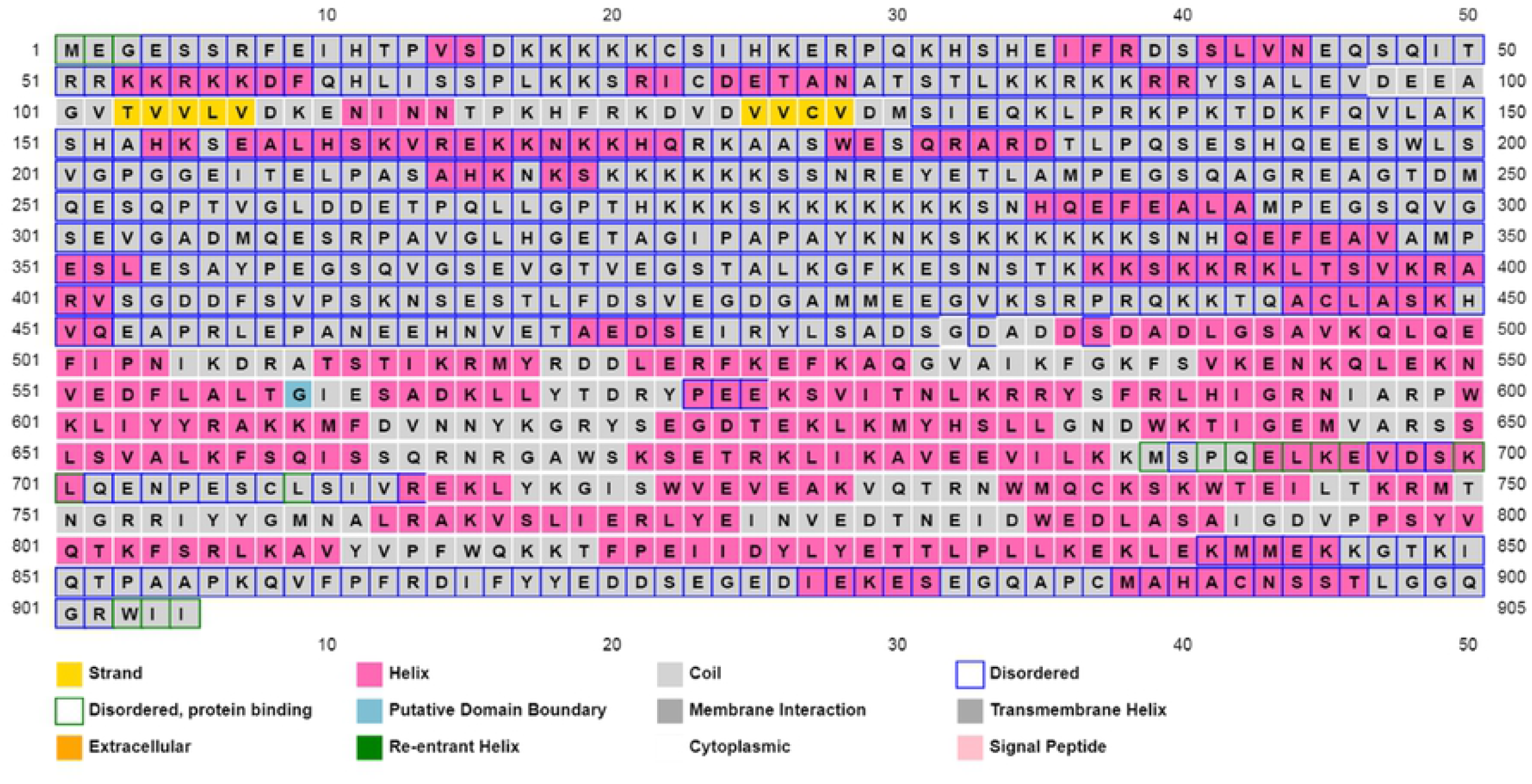
Sequence-based representation of the disordered and protein binding regions of TTF1, as predicted using PsiPred.

**S 2 Fig.**
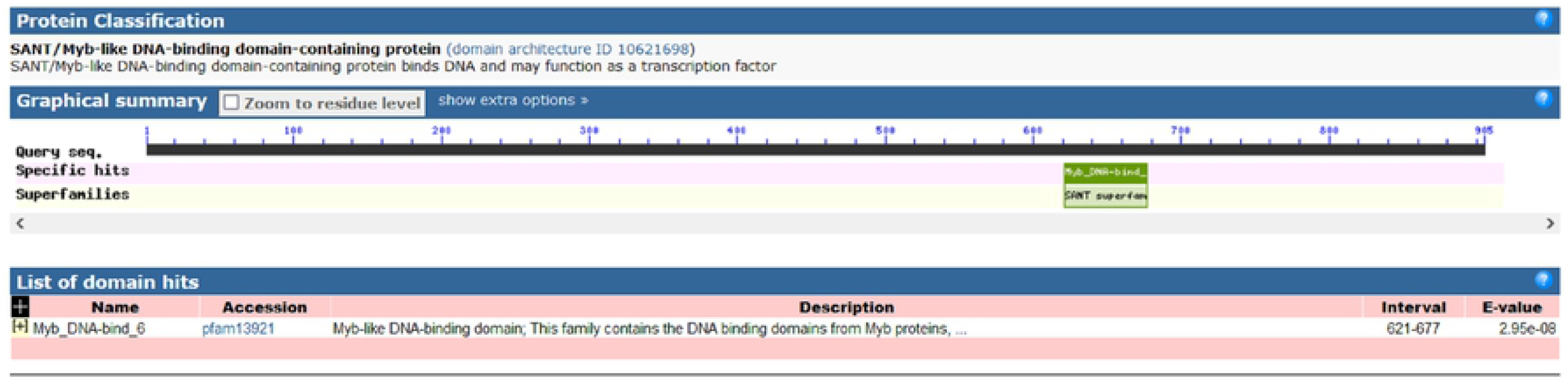
Results of CD search indicating the presence of a SANT/Myb-like DNA-binding domain in TTF1.

**S 1 Table.** Occupancy of the hydrogen bonds formed by the central residues throughout the trajectory and in the last 50 ns.

| Donor | Acceptor | Occupancy (200 ns) | Occupancy (last 50 ns) |
| --- | --- | --- | --- |
| K68*-Main | L67-Side | 75.34% | 85.41% |
| A175-Main | A176*-Main | 76.84% | 78.12% |
| R71*-Main | K68*-Main | 77.29% | 83.02% |
| S197-Side | E195*-Side | 77.46% | 99.50% |
| E27*-Main | I24-Main | 77.51% | 81.02% |
| D471*-Main | A469-Main | 77.66% | 97.60% |
| T896*-Main | N893*-Main | 78.04% | 97.60% |
| E882*-Main | E880-Side | 78.41% | 94.21% |
| D185-Side | N228*-Side | 78.66% | 82.72% |
| S227-Side | R229*-Side | 79.01% | 119.18% |
| K17*-Main | D16-Side | 79.29% | 91.01% |
| R182*-Side | N228*-Side | 79.36% | 80.92% |
| E431*-Main | A427-Main | 79.41% | 91.31% |
| E209-Main | E206*-Main | 79.41% | 93.61% |
| P437-Side | K434*-Main | 79.76% | 88.81% |
| W198*-Side | G202*-Main | 79.94% | 81.72% |
| I474-Main | D471*-Main | 79.99% | 90.11% |
| S197-Main | W198*-Main | 80.58% | 83.12% |
| A183-Main | R182*-Side | 81.46% | 75.82% |
| F657*-Main | K656-Side | 82.58% | 87.91% |
| E431*-Side | A427-Main | 82.83% | 117.98% |
| K18-Main | K17*-Side | 83.03% | 99.00% |
| P203-Side | W198*-Main | 83.56% | 125.27% |
| E35*-Main | H34*-Side | 84.43% | 82.82% |
| K31*-Side | E27*-Side | 84.46% | 125.47% |
| F657*-Main | V653-Main | 84.53% | 82.52% |
| R164*-Main | V163-Side | 85.01% | 94.91% |
| K31*-Main | H32-Main | 85.61% | 86.21% |
| E430-Main | E431*-Main | 85.63% | 90.91% |
| A176*-Main | S177-Main | 86.01% | 82.12% |
| S23*-Main | K19-Main | 86.01% | 100.00% |
| Q30*-Side | E27*-Main | 86.23% | 87.21% |
| S895*-Main | N893*-Side | 86.26% | 95.40% |
| K656-Main | F657*-Main | 86.38% | 86.81% |
| D471*-Main | S472-Main | 86.66% | 92.11% |
| V433-Main | K434*-Main | 87.06% | 82.32% |
| F657*-Side | V653-Main | 87.33% | 95.70% |
| K434*-Main | S435*-Main | 87.43% | 94.61% |
| S661-Side | F657*-Main | 87.91% | 78.82% |
| S33-Main | Q30*-Main | 88.28% | 90.11% |
| E27*-Main | R28-Main | 88.51% | 88.31% |
| A176*-Main | Q172*-Main | 88.93% | 95.00% |
| K31*-Side | E27*-Main | 89.63% | 85.31% |
| R438-Main | S435*-Main | 89.68% | 94.11% |
| D471*-Main | E467-Main | 89.81% | 90.51% |
| I660*-Main | F657*-Main | 89.93% | 93.61% |
| Q30*-Main | E27*-Main | 90.08% | 92.71% |
| P188-Side | N228*-Main | 90.83% | 115.78% |
| E431*-Main | G432-Main | 91.03% | 97.20% |
| S435*-Main | E431*-Main | 91.05% | 94.61% |
| R664*-Main | I660*-Main | 91.45% | 126.17% |
| E882*-Main | S881-Side | 91.93% | 94.71% |
| E35*-Main | I36-Main | 92.25% | 88.01% |
| Q181-Side | R182*-Main | 92.65% | 87.61% |
| T896*-Main | S895*-Side | 92.85% | 81.02% |
| K68*-Main | S65-Main | 93.45% | 84.82% |
| S435*-Main | K434*-Side | 93.58% | 96.20% |
| R229*-Main | N228*-Side | 93.73% | 97.80% |
| S661-Main | F657*-Main | 93.83% | 89.11% |
| I660*-Main | S661-Main | 93.85% | 98.60% |
| Q439-Main | S435*-Main | 93.88% | 92.51% |
| G901-Main | G899*-Main | 94.28% | 99.50% |
| E27*-Main | S23*-Main | 94.65% | 94.31% |
| A176*-Main | A175-Side | 94.73% | 97.70% |
| R71*-Main | S70-Side | 95.15% | 114.89% |
| S472-Main | D471*-Side | 95.33% | 93.91% |
| Q30*-Main | K31*-Main | 95.33% | 96.20% |
| I36-Main | E35*-Side | 95.38% | 92.11% |
| L62-Side | H34*-Main | 95.68% | 112.79% |
| E35*-Main | H32-Main | 96.10% | 91.71% |
| R436-Main | K434*-Main | 96.93% | 93.11% |
| S190-Main | E191*-Main | 97.15% | 97.20% |
| E470-Main | D471*-Main | 97.18% | 95.40% |
| E27*-Main | K26-Side | 97.50% | 96.80% |
| K26-Main | S23*-Main | 97.68% | 98.20% |
| H34*-Main | E35*-Main | 97.73% | 95.90% |
| I36-Main | H34*-Main | 98.10% | 123.08% |
| G432-Main | E431*-Side | 98.18% | 98.50% |
| K166-Main | E165*-Side | 98.18% | 99.30% |
| S435*-Main | R436-Main | 98.25% | 96.60% |
| D471*-Main | E470-Side | 98.43% | 97.00% |
| R173-Main | Q172*-Side | 98.45% | 104.40% |
| L67-Main | K68*-Main | 98.55% | 96.60% |
| S200-Main | E196*-Main | 98.70% | 96.50% |
| N228*-Main | T186*-Main | 99.08% | 99.90% |
| L199-Main | E195*-Main | 99.18% | 99.40% |
| T896*-Main | L897-Main | 99.30% | 96.60% |
| W198*-Main | E195*-Main | 99.30% | 99.50% |
| E195*-Main | Q194-Side | 99.50% | 99.40% |
| V201-Main | W198*-Main | 99.55% | 99.10% |
| W903-Main | R902*-Side | 99.75% | 100.30% |
| W198*-Main | L199-Main | 99.80% | 100.00% |
| K31*-Main | E27*-Main | 99.85% | 99.60% |
| T186*-Main | D185-Side | 99.90% | 99.90% |
| E195*-Main | K700-Main | 99.98% | 99.70% |
| D185-Main | T186*-Main | 100.00% | 100.00% |
| E165*-Main | K166-Main | 100.02% | 100.00% |
| T186*-Main | L187-Main | 100.02% | 100.10% |
| E195*-Main | E196*-Main | 100.05% | 100.10% |
| E230-Main | R229*-Side | 100.07% | 96.80% |
| Q194-Main | E195*-Main | 100.07% | 100.00% |
| H160-Main | S161*-Main | 100.07% | 100.10% |
| C892-Main | N893*-Main | 100.22% | 100.20% |
| R164*-Main | E165*-Main | 100.27% | 100.20% |
| R71*-Main | I72-Main | 100.55% | 101.20% |
| K26-Main | E27*-Main | 100.57% | 100.40% |
| S161*-Main | K162-Main | 101.10% | 100.10% |
| K31*-Main | Q30*-Side | 101.15% | 101.80% |
| C22-Main | S23*-Main | 101.20% | 105.19% |
| R229*-Main | E230-Main | 101.22% | 101.90% |
| E95-Main | L94-Side | 101.35% | 104.10% |
| D39-Main | R38*-Side | 101.55% | 102.80% |
| Q30*-Side | K31*-Main | 101.65% | 102.90% |
| F37-Main | R38*-Main | 101.97% | 114.59% |
| S227-Side | N228*-Main | 102.00% | 97.30% |
| N228*-Main | R229*-Main | 102.32% | 100.20% |
| S23*-Main | C22-Side | 102.35% | 94.41% |
| R436-Main | S435*-Side | 102.40% | 94.01% |
| V163-Side | R164*-Main | 102.45% | 96.60% |
| V201-Main | G202*-Main | 102.62% | 103.70% |
| K17*-Main | K18-Main | 102.80% | 96.40% |
| E430-Side | E431*-Main | 102.85% | 93.61% |
| R182*-Side | A176*-Main | 102.90% | 130.57% |
| K166-Main | E165*-Main | 103.42% | 102.20% |
| S192-Main | E191*-Side | 103.70% | 96.40% |
| N893*-Main | S894-Main | 104.27% | 104.10% |
| S33-Side | Q30*-Main | 104.37% | 95.00% |
| K68*-Main | K69-Main | 104.37% | 98.50% |
| F657*-Main | S658-Main | 104.40% | 100.70% |
| L67-Side | K68*-Main | 104.57% | 144.26% |
| S70-Main | R71*-Main | 104.70% | 95.10% |
| E191*-Main | S190-Side | 105.00% | 102.60% |
| G899*-Main | L897-Main | 105.77% | 158.34% |
| H32-Main | K31*-Side | 105.95% | 100.10% |
| H171-Main | Q172*-Main | 106.65% | 101.40% |
| L897-Main | S895*-Main | 106.80% | 133.17% |
| S895*-Side | E165*-Side | 106.85% | 204.70% |
| Q663-Main | R664*-Main | 107.02% | 117.68% |
| S881-Side | E882*-Main | 107.50% | 138.56% |
| E35*-Main | S33-Main | 107.55% | 100.40% |
| E431*-Main | E430-Side | 107.60% | 113.99% |
| R902*-Main | W903-Main | 107.62% | 104.10% |
| K225-Main | S226*-Main | 107.67% | 107.29% |
| S161*-Main | H160-Side | 109.75% | 95.40% |
| R28-Main | E27*-Side | 110.17% | 107.49% |
| S881-Main | E882*-Main | 111.24% | 105.79% |
| E882*-Main | G883-Main | 111.49% | 115.68% |
| K434*-Main | V433-Side | 111.57% | 127.37% |
| K434*-Side | Q663-Main | 111.84% | 118.18% |
| S227-Main | N228*-Main | 113.14% | 115.48% |
| Q30*-Main | P29-Side | 113.47% | 113.29% |
| S435*-Main | V433-Main | 114.04% | 104.90% |
| S226*-Main | K224-Main | 114.97% | 136.76% |
| K174-Main | Q172*-Main | 115.02% | 127.67% |
| K879-Side | R902*-Side | 115.34% | 217.68% |
| S895*-Side | T896*-Main | 115.49% | 80.92% |
| S661-Main | I660*-Side | 115.52% | 124.58% |
| R902*-Main | Q900-Main | 116.44% | 129.77% |
| I660*-Main | Q659-Side | 116.72% | 140.86% |
| E165*-Main | R164*-Side | 117.87% | 159.94% |
| S40-Main | R38*-Main | 117.89% | 130.77% |
| K434*-Main | G432-Main | 118.02% | 156.04% |
| V433-Main | E431*-Main | 118.04% | 109.99% |
| R38*-Side | E35*-Main | 118.32% | 105.09% |
| N893*-Side | R164*-Side | 118.77% | 215.38% |
| F37-Main | E35*-Main | 118.79% | 126.77% |
| W198*-Side | L199-Main | 119.27% | 118.88% |
| A176*-Side | Q172*-Main | 119.49% | 89.71% |
| F657*-Side | A454-Main | 119.77% | 171.13% |
| S435*-Side | R436-Main | 120.59% | 144.56% |
| R902*-Side | K879-Side | 120.66% | 247.15% |
| V433-Side | K434*-Main | 120.76% | 134.97% |
| R184-Main | R182*-Main | 120.99% | 114.99% |
| H34*-Main | S33-Side | 122.36% | 111.69% |
| E431*-Main | M429-Main | 122.46% | 117.48% |
| S70-Side | R71*-Main | 122.59% | 123.88% |
| T896*-Side | N893*-Main | 123.06% | 208.99% |
| K68*-Main | P66-Main | 126.81% | 99.40% |
| P29-Side | Q30*-Main | 127.76% | 125.87% |
| I660*-Side | S661-Main | 127.76% | 135.36% |
| R164*-Side | N893*-Side | 128.19% | 248.15% |
| I660*-Main | S658-Main | 128.79% | 173.53% |
| Q172*-Side | K803-Main | 129.04% | 122.78% |
| R38*-Main | I36-Main | 129.06% | 104.90% |
| K68*-Side | K69-Main | 129.29% | 105.89% |
| S33-Main | K31*-Main | 131.36% | 139.16% |
| S662-Main | I660*-Main | 131.58% | 86.01% |
| G202*-Main | W198*-Side | 134.06% | 118.28% |
| E206*-Main | G204-Main | 134.06% | 186.01% |
| N893*-Main | C892-Side | 134.31% | 126.77% |
| G205-Main | E206*-Main | 134.93% | 118.18% |
| E35*-Side | I36-Main | 135.38% | 125.47% |
| E191*-Side | S192-Main | 135.86% | 145.65% |
| K17*-Side | K18-Main | 136.01% | 134.17% |
| V163-Main | R164*-Main | 136.38% | 148.45% |
| R38*-Side | D39-Main | 136.88% | 107.79% |
| E196*-Side | S197-Main | 137.48% | 136.06% |
| H160-Side | S161*-Main | 137.56% | 185.51% |
| R182*-Side | A183-Main | 137.63% | 136.76% |
| C22-Side | S23*-Main | 137.98% | 153.55% |
| S197-Main | E196*-Side | 138.01% | 107.19% |
| R71*-Side | I72-Main | 138.53% | 132.07% |
| R664*-Main | N665-Main | 138.96% | 156.54% |
| S197-Side | W198*-Main | 139.06% | 130.97% |
| G202*-Main | W198*-Main | 139.33% | 141.26% |
| A176*-Side | S177-Main | 139.86% | 141.46% |
| W198*-Side | S699-Side | 141.23% | 190.21% |
| E431*-Side | G432-Main | 141.73% | 172.33% |
| S895*-Side | N893*-Side | 141.73% | 173.23% |
| S894-Side | S895*-Main | 143.00% | 136.36% |
| D16-Side | K17*-Main | 143.30% | 147.75% |
| K656-Side | F657*-Main | 143.50% | 157.84% |
| E473-Main | D471*-Main | 143.68% | 156.84% |
| S895*-Main | S894-Side | 145.23% | 128.37% |
| F657*-Main | L655-Main | 145.63% | 145.45% |
| R229*-Side | S227-Side | 145.80% | 215.18% |
| S161*-Side | K162-Main | 146.63% | 154.05% |
| Q663-Side | R664*-Main | 146.88% | 101.10% |
| I24-Main | S23*-Side | 146.90% | 160.14% |
| S435*-Side | E431*-Main | 146.98% | 139.66% |
| S23*-Side | I24-Main | 148.58% | 131.57% |
| T896*-Side | L897-Main | 148.63% | 178.42% |
| H32-Main | Q30*-Main | 148.75% | 144.16% |
| K434*-Side | S435*-Main | 149.48% | 165.43% |
| S894-Main | N893*-Side | 150.75% | 151.95% |
| W198*-Main | S197-Side | 151.25% | 157.04% |
| L897-Main | T896*-Side | 152.00% | 182.52% |
| Q194-Side | E195*-Main | 152.55% | 155.44% |
| R902*-Side | T896*-Main | 152.85% | 236.46% |
| A176*-Main | K174-Main | 153.85% | 155.94% |
| D471*-Side | S472-Main | 154.42% | 129.07% |
| E470-Side | D471*-Main | 154.82% | 117.38% |
| K167-Main | E165*-Main | 154.90% | 170.13% |
| Q659-Side | I660*-Main | 156.85% | 186.51% |
| G202*-Main | V201-Side | 157.22% | 152.05% |
| K31*-Side | H32-Main | 157.72% | 183.82% |
| G898-Main | G899*-Main | 158.45% | 136.66% |
| Q30*-Main | R28-Main | 159.10% | 163.44% |
| L199-Main | W198*-Side | 159.12% | 164.04% |
| S33-Side | H34*-Main | 159.62% | 156.54% |
| E206*-Side | I207-Main | 159.67% | 127.57% |
| V201-Side | G202*-Main | 159.82% | 169.03% |
| H193-Main | E191*-Main | 160.04% | 161.24% |
| T208-Main | E206*-Main | 160.74% | 192.31% |
| S70-Main | K68*-Main | 161.44% | 179.72% |
| H34*-Main | H32-Main | 163.47% | 161.64% |
| Q30*-Side | R71*-Main | 165.09% | 194.81% |
| R164*-Side | V163-Main | 166.04% | 191.01% |
| W198*-Main | E196*-Main | 167.82% | 178.82% |
| K162-Main | S161*-Side | 168.42% | 177.02% |
| K17*-Main | S15-Main | 168.49% | 147.05% |
| C892-Side | N893*-Main | 168.52% | 151.65% |
| H34*-Side | E35*-Main | 168.84% | 165.63% |
| D185-Side | T186*-Main | 169.34% | 169.83% |
| Q659-Main | F657*-Main | 171.14% | 183.42% |
| S190-Side | E191*-Main | 171.89% | 188.21% |
| S226*-Main | S227-Main | 171.91% | 160.04% |
| K391-Side | E431*-Side | 173.11% | 525.57% |
| Q172*-Main | H171-Side | 174.26% | 173.13% |
| K26-Side | E27*-Main | 174.71% | 170.13% |
| W178-Main | A176*-Main | 174.79% | 172.33% |
| S200-Side | E196*-Main | 175.24% | 167.83% |
| T896*-Main | S894-Main | 175.56% | 184.32% |
| G901-Main | R902*-Main | 175.79% | 185.51% |
| E27*-Side | R28-Main | 176.59% | 143.36% |
| R664*-Side | E453-Side | 177.26% | 163.94% |
| R229*-Side | E230-Main | 178.89% | 193.61% |
| E195*-Side | E196*-Main | 181.06% | 179.82% |
| N228*-Main | S227-Side | 181.73% | 188.41% |
| R164*-Side | E882*-Side | 182.91% | 338.56% |
| K434*-Side | E430-Main | 183.93% | 255.14% |
| S895*-Main | N893*-Main | 184.93% | 197.50% |
| N228*-Side | R229*-Main | 185.01% | 193.71% |
| L187-Main | T186*-Side | 185.56% | 190.91% |
| R902*-Side | W903-Main | 186.46% | 191.41% |
| Q172*-Side | R173-Main | 186.56% | 181.12% |
| R164*-Side | E165*-Main | 187.01% | 173.33% |
| A175-Side | A176*-Main | 188.11% | 193.61% |
| H25-Main | S23*-Main | 189.38% | 194.61% |
| W198*-Side | Q194-Side | 190.18% | 201.00% |
| G202*-Main | P203-Main | 190.80% | 196.10% |
| K17*-Side | E206*-Side | 192.30% | 445.15% |
| S197-Main | E195*-Main | 193.05% | 194.41% |
| G899*-Main | Q900-Main | 194.33% | 200.00% |
| E165*-Side | K166-Main | 194.58% | 198.40% |
| R182*-Main | S180-Main | 194.80% | 195.40% |
| N228*-Side | T186*-Main | 197.88% | 215.28% |
| S200-Main | W198*-Main | 199.08% | 198.20% |
| G202*-Main | P203-Side | 199.68% | 200.00% |
| P203-Main | G202*-Main | 200.00% | 200.00% |
| S226*-Main | K225-Main | 200.12% | 200.20% |
| R902*-Side | G901-Main | 204.92% | 249.75% |
| S649-Side | D471*-Side | 208.30% | 151.15% |
| K166-Side | E165*-Main | 214.44% | 215.08% |
| S699-Side | W198*-Side | 215.84% | 262.04% |
| R664*-Main | Q663-Main | 217.54% | 200.20% |
| K434*-Side | E431*-Side | 222.09% | 399.30% |
| K17*-Side | P13-Main | 224.49% | 154.05% |
| S226*-Side | K225-Main | 227.46% | 213.69% |
| K69-Side | E35*-Side | 229.14% | 129.47% |
| E196*-Main | E195*-Side | 237.91% | 280.82% |
| K17*-Side | P203-Main | 239.81% | 142.26% |
| N228*-Side | D185-Main | 240.05% | 254.15% |
| R664*-Side | Q663-Main | 248.33% | 260.14% |
| N615-Side | D471*-Side | 252.77% | 228.37% |
| K17*-Side | G204-Main | 275.61% | 254.55% |
| S227-Side | E191*-Side | 276.89% | 290.61% |
| K17*-Side | V201-Main | 277.41% | 122.28% |
| S227-Side | S226*-Main | 278.39% | 270.13% |
| R71*-Side | E9-Side | 284.26% | 79.12% |
| K434*-Side | E430-Side | 288.83% | 660.24% |
| K17*-Side | G202*-Main | 295.48% | 306.79% |
| T186*-Main | D185-Main | 300.47% | 300.50% |
| E195*-Main | Q194-Main | 301.10% | 301.20% |
| E165*-Main | R164*-Main | 301.17% | 300.40% |
| E196*-Main | E195*-Main | 301.75% | 300.40% |
| N893*-Main | C892-Main | 304.20% | 309.79% |
| L187-Main | T186*-Main | 304.65% | 302.40% |
| R229*-Main | N228*-Main | 305.92% | 302.10% |
| S161*-Main | H160-Main | 306.82% | 306.69% |
| W903-Main | R902*-Main | 308.55% | 304.20% |
| K162-Main | S161*-Main | 312.37% | 312.69% |
| Q900-Main | G899*-Main | 312.69% | 300.00% |
| N228*-Main | S227-Main | 313.14% | 315.48% |
| R164*-Side | E880-Side | 313.72% | 552.85% |
| Q172*-Main | H171-Main | 336.23% | 334.97% |
| I72-Main | R71*-Main | 338.01% | 341.26% |
| E206*-Main | G205-Main | 340.43% | 400.00% |
| S894-Main | N893*-Main | 349.65% | 346.95% |
| T186*-Side | L187-Main | 359.20% | 397.00% |
| R164*-Main | V163-Main | 366.74% | 361.24% |
| R902*-Side | E869-Side | 380.26% | 562.74% |
| R902*-Main | G901-Main | 384.53% | 400.00% |
| R229*-Side | E191*-Side | 388.03% | 500.00% |
| R182*-Side | E232-Main | 389.63% | 413.49% |
| N665-Main | R664*-Main | 392.23% | 400.00% |
| G432-Main | E431*-Main | 399.93% | 399.90% |
| R38*-Main | F37-Main | 399.95% | 400.00% |
| L897-Main | T896*-Main | 400.00% | 400.00% |
| T896*-Main | S895*-Main | 400.00% | 400.00% |
| S895*-Main | S894-Main | 400.00% | 400.00% |
| E882*-Main | S881-Main | 400.00% | 400.00% |
| R173-Main | Q172*-Main | 400.00% | 400.00% |
| E27*-Main | K26-Main | 400.00% | 400.00% |
| Q30*-Main | P29-Main | 400.00% | 400.00% |
| R28-Main | E27*-Main | 400.00% | 400.00% |
| S23*-Main | C22-Main | 400.00% | 400.00% |
| I24-Main | S23*-Main | 400.00% | 400.00% |
| H34*-Main | S33-Main | 400.00% | 400.00% |
| E35*-Main | H34*-Main | 400.00% | 400.00% |
| K31*-Main | Q30*-Main | 400.00% | 400.00% |
| H32-Main | K31*-Main | 400.00% | 400.00% |
| I36-Main | E35*-Main | 400.00% | 400.00% |
| A176*-Main | A175-Main | 400.00% | 400.00% |
| S177-Main | A176*-Main | 400.00% | 400.00% |
| K69-Main | K68*-Main | 400.00% | 400.00% |
| R71*-Main | S70-Main | 400.00% | 400.00% |
| S227-Main | S226*-Main | 400.00% | 400.00% |
| I207-Main | E206*-Main | 400.00% | 400.00% |
| K17*-Main | D16-Main | 400.00% | 400.00% |
| K18-Main | K17*-Main | 400.00% | 400.00% |
| K68*-Main | L67-Main | 400.00% | 400.00% |
| D39-Main | R38*-Main | 400.00% | 400.00% |
| E230-Main | R229*-Main | 400.00% | 400.00% |
| R182*-Main | Q181-Main | 400.00% | 400.00% |
| A183-Main | R182*-Main | 400.00% | 400.00% |
| L199-Main | W198*-Main | 400.00% | 400.00% |
| S192-Main | E191*-Main | 400.00% | 400.00% |
| E191*-Main | S190-Main | 400.00% | 400.00% |
| W198*-Main | S197-Main | 400.00% | 400.00% |
| S197-Main | E196*-Main | 400.00% | 400.00% |
| E431*-Main | E430-Main | 400.00% | 400.00% |
| R436-Main | S435*-Main | 400.00% | 400.00% |
| S435*-Main | K434*-Main | 400.00% | 400.00% |
| K434*-Main | V433-Main | 400.00% | 400.00% |
| S658-Main | F657*-Main | 400.00% | 400.00% |
| I660*-Main | Q659-Main | 400.00% | 400.00% |
| F657*-Main | K656-Main | 400.00% | 400.00% |
| S661-Main | I660*-Main | 400.00% | 400.00% |
| D471*-Main | E470-Main | 400.00% | 400.00% |
| S472-Main | D471*-Main | 400.00% | 400.00% |
| G899*-Main | G898-Main | 400.10% | 401.00% |
| G202*-Main | V201-Main | 400.82% | 403.30% |
| R38*-Side | E35*-Side | 406.17% | 385.11% |
| R229*-Side | N228*-Main | 421.29% | 512.19% |
| R164*-Side | E165*-Side | 457.75% | 489.11% |
| G883-Main | E882*-Main | 472.74% | 561.44% |
| K20-Side | E27*-Side | 530.31% | 389.51% |
| P203-Side | G202*-Main | 601.07% | 602.20% |
| K765-Side | E206*-Side | 655.37% | 665.53% |

The central residues are marked by \*. Side and Main indicate whether the hydrogen bond was formed with an atom of the side chain or the main chain, respectively. Occupancies >100% indicate the formation of more than one inter-residue hydrogen bond.

## References

1. Zhou H, Wang Y, Lv Q, Zhang J, Wang Q, Gao F, et al. Overexpression of ribosomal RNA in the development of human cervical cancer is associated with rDNA promoter hypomethylation. PLoS One. 2016;11: e0163340. doi:10.1371/journal.pone.0163340

2. Akamatsu Y, Kobayashi T. The Human RNA Polymerase I Transcription Terminator Complex Acts as a Replication Fork Barrier That Coordinates the Progress of Replication with rRNA Transcription Activity. Mol Cell Biol. 2015;35: 1871–1881. doi:10.1128/mcb.01521-14

3. Evers R, Grummt I. Molecular coevolution of mammalian ribosomal gene terminator sequences and the transcription termination factor TTF-I. Proc Natl Acad Sci U S A. 1995;92: 5827–5831. doi:10.1073/pnas.92.13.5827

4. Pütter V, Grummt F. Transcription termination factor TTF-I exhibits contrahelicase activity during DNA replication. EMBO Rep. 2002;3: 147–152. doi:10.1093/embo-reports/kvf027

5. Längst G, Blank TA, Becker PB, Grummt I. RNA polymerase I transcription on nucleosomal templates: The transcription termination factor TTF-I induces chromatin remodeling and relieves transcriptional repression. EMBO J. 1997;16: 760–768. doi:10.1093/emboj/16.4.760

6. Aamann MD, Muftuoglu M, Bohr VA, Stevnsner T. Multiple interaction partners for Cockayne syndrome proteins: Implications for genome and transcriptome maintenance. Mech Ageing Dev. 2013;134: 212–224. doi:10.1016/j.mad.2013.03.009

7. Lessard F, Stefanovsky V, Tremblay MG, Moss T. The cellular abundance of the essential transcription termination factor TTF-I regulates ribosome biogenesis and is determined by MDM2 ubiquitinylation. Nucleic Acids Res. 2012;40: 5357–5367. doi:10.1093/nar/gks198

8. Lessard F, Morin F, Ivanchuk S, Langlois F, Stefanovsky V, Rutka J, et al. The ARF Tumor Suppressor Controls Ribosome Biogenesis by Regulating the RNA Polymerase I Transcription Factor TTF-I. Mol Cell. 2010;38: 539–550. doi:10.1016/j.molcel.2010.03.015

9. Stults DM, Killen MW, Williamson EP, Hourigan JS, Vargas HD, Arnold SM, et al. Human rRNA gene clusters are recombinational hotspots in cancer. Cancer Res. 2009;69: 9096–9104. doi:10.1158/0008-5472.CAN-09-2680

10. Komatsu H, Iguchi T, Ueda M, Nambara S, Saito T, Hirata H, et al. Clinical and biological significance of transcription termination factor, RNA polymerase i in human liver hepatocellular carcinoma. Oncol Rep. 2016;35: 2073–2080. doi:10.3892/or.2016.4593

11. Ueda M, Iguchi T, Nambara S, Saito T, Komatsu H, Sakimura S, et al. Overexpression of Transcription Termination Factor 1 is Associated with a Poor Prognosis in Patients with Colorectal Cancer. Ann Surg Oncol. 2015;22: 1490–1498. doi:10.1245/s10434-015-4652-7

12. Sander EE, Grummt I. Oligomerization of the transcription termination factor TTF-I: Implications for the structural organization of ribosomal transcription units. Nucleic Acids Res. 1997;25: 1142–1147. doi:10.1093/nar/25.6.1142

13. Németh A, Guibert S, Tiwari VK, Ohlsson R, Längst G. Epigenetic regulation of TTF-I-mediated promoter-terminator interactions of rRNA genes. EMBO J. 2008;27: 1255–1265. doi:10.1038/emboj.2008.57

14. Boutin J, Lessard F, Tremblay MG, Moss T. The short N-terminal repeats of transcription termination factor 1 contain semi-redundant nucleolar localization signals and P19-ARF tumor suppressor binding sites. Yale J Biol Med. 2019;92: 385–396. Available: /pmc/articles/PMC6747939/

15. Jaiswal R, Choudhury M, Zaman S, Singh S, Santosh V, Bastia D, et al. Functional architecture of the Reb1-Ter complex of Schizosaccharomyces pombe. Proc Natl Acad Sci U S A. 2016;113: E2267–E2276. doi:10.1073/pnas.1525465113

16. Park SH, Yu KL, Jung YM, Lee SD, Kim MJ, You JC. Investigation of functional roles of transcription termination factor-1 (TTF-I) in HIV-1 replication. BMB Rep. 2018;51: 338–343. doi:10.5483/BMBRep.2018.51.7.032

17. Singh SK, Sabatinos S, Forsburg S, Bastia D. Regulation of Replication Termination by Reb1 Protein-Mediated Action at a Distance. Cell. 2010;142: 868–878. doi:10.1016/j.cell.2010.08.013

18. Khor BY, Tye GJ, Lim TS, Choong YS. General overview on structure prediction of twilight-zone proteins. Theor Biol Med Model. 2015;12. doi:10.1186/s12976-015-0014-1

19. Chung SY, Subbiah S. A structural explanation for the twilight zone of protein sequence homology. Structure. 1996;4: 1123–1127. doi:10.1016/S0969-2126(96)00119-0

20. Roy A, Kucukural A, Zhang Y. I-TASSER: A unified platform for automated protein structure and function prediction. Nat Protoc. 2010;5: 725–738. doi:10.1038/nprot.2010.5

21. Yang J, Yan R, Roy A, Xu D, Poisson J, Zhang Y. The I-TASSER suite: Protein structure and function prediction. Nature Methods. Nat Methods; 2014. pp. 7–8. doi:10.1038/nmeth.3213

22. Kryshtafovych A, Schwede T, Topf M, Fidelis K, Moult J. Critical assessment of methods of protein structure prediction (CASP)—Round XIII. Proteins: Structure, Function and Bioinformatics. Proteins; 2019. pp. 1011–1020. doi:10.1002/prot.25823

23. Rost B. Twilight zone of protein sequence alignments. Protein Eng. 1999;12: 85–94. doi:10.1093/protein/12.2.85

24. Krieger E, Joo K, Lee J, Lee J, Raman S, Thompson J, et al. Improving physical realism, stereochemistry, and side-chain accuracy in homology modeling: Four approaches that performed well in CASP8. Proteins: Structure, Function and Bioinformatics. Proteins; 2009. pp. 114–122. doi:10.1002/prot.22570

25. M W, MJ S. ProSA-web: interactive web service for the recognition of errors in three-dimensional structures of proteins. Nucleic Acids Res. 2007;35. doi:10.1093/NAR/GKM290

26. Zhang B, Ye W, Ye Y, Zhou H, Saeed AFUH, Chen J, et al. Structural insights into Cas13b-guided CRISPR RNA maturation and recognition. Cell Research. Cell Res; 2018. pp. 1198–1201. doi:10.1038/s41422-018-0109-4

27. Jumper J, Evans R, Pritzel A, Green T, Figurnov M, Ronneberger O, et al. Highly accurate protein structure prediction with AlphaFold. Nature. 2021;596: 583–589. doi:10.1038/s41586-021-03819-2

28. Gasteiger E, Hoogland C, Gattiker A, Duvaud S, Wilkins MR, Appel RD, et al. Protein Identification and Analysis Tools on the ExPASy Server. The Proteomics Protocols Handbook. Humana Press; 2005. pp. 571–607. doi:10.1385/1-59259-890-0:571

29. Jones DT, Cozzetto D. DISOPRED3: Precise disordered region predictions with annotated protein-binding activity. Bioinformatics. 2015;31: 857–863. doi:10.1093/bioinformatics/btu744

30. Ward JJ, McGuffin LJ, Bryson K, Buxton BF, Jones DT. The DISOPRED server for the prediction of protein disorder. Bioinformatics. 2004;20: 2138–2139. doi:10.1093/bioinformatics/bth195

31. Wiederstein M, Sippl MJ. ProSA-web: Interactive web service for the recognition of errors in three-dimensional structures of proteins. Nucleic Acids Res. 2007;35: W407–10. doi:10.1093/nar/gkm290

32. Laskowski RA, MacArthur MW, Moss DS, Thornton JM. PROCHECK: a program to check the stereochemical quality of protein structures. J Appl Crystallogr. 1993;26: 283–291. doi:10.1107/s0021889892009944

33. Zhang Y, Skolnick J. TM-align: A protein structure alignment algorithm based on the TM-score. Nucleic Acids Res. 2005;33: 2302–2309. doi:10.1093/nar/gki524

34. Zhang Y, Skolnick J. Scoring function for automated assessment of protein structure template quality. Proteins Struct Funct Genet. 2004;57: 702–710. doi:10.1002/prot.20264

35. Yang J, Roy A, Zhang Y. Protein-ligand binding site recognition using complementary binding-specific substructure comparison and sequence profile alignment. Bioinformatics. 2013;29: 2588–2595. doi:10.1093/bioinformatics/btt447

36. Zhang C, Freddolino PL, Zhang Y. COFACTOR: Improved protein function prediction by combining structure, sequence and protein-protein interaction information. Nucleic Acids Res. 2017;45: W291–W299. doi:10.1093/nar/gkx366

37. Lu S, Wang J, Chitsaz F, Derbyshire MK, Geer RC, Gonzales NR, et al. CDD/SPARCLE: The conserved domain database in 2020. Nucleic Acids Res. 2020;48: D265–D268. doi:10.1093/nar/gkz991

38. Humphrey W, Dalke A, Schulten K. VMD: Visual molecular dynamics. J Mol Graph. 1996;14: 33–38. doi:10.1016/0263-7855(96)00018-5

39. Knapp B, Lederer N, Omasits U, Schreiner W. VmdICE: A plug-in for rapid evaluation of molecular dynamics simulations using VMD. J Comput Chem. 2010;31: 2868–2873. doi:10.1002/jcc.21581

40. Pettersen EF, Goddard TD, Huang CC, Couch GS, Greenblatt DM, Meng EC, et al. UCSF Chimera - A visualization system for exploratory research and analysis. J Comput Chem. 2004;25: 1605–1612. doi:10.1002/jcc.20084

41. Shannon P, Markiel A, Ozier O, Baliga NS, Wang JT, Ramage D, et al. Cytoscape: A software Environment for integrated models of biomolecular interaction networks. Genome Res. 2003;13: 2498–2504. doi:10.1101/gr.1239303

42. Doncheva NT, Klein K, Domingues FS, Albrecht M. Analyzing and visualizing residue networks of protein structures. Trends in Biochemical Sciences. Trends Biochem Sci; 2011. pp. 179–182. doi:10.1016/j.tibs.2011.01.002

